# Pervasive additive and non-additive effects within the HLA region contribute to disease risk in the UK Biobank

**DOI:** 10.1101/2020.05.28.119669

**Authors:** Guhan Ram Venkataraman, Julia Eve Olivieri, Christopher DeBoever, Yosuke Tanigawa, Johanne Marie Justesen, Alexander Dilthey, Manuel A. Rivas

## Abstract

The human leukocyte antigen (HLA) region is one of the most disease-associated regions of the human genome, yet even well-studied alleles in the HLA region have unknown impact on disease. Here, we study the effect of 156 HLA alleles on 677 binary phenotypes for 337,138 individuals in the UK Biobank. We assess HLA allele associations and subsequently use Bayesian Model Averaging for conditional analysis, a) replicating 88 known associations between HLA alleles and binary disease phenotypes such as cancer, and b) discovering 90 novel associations to phenotypes such as skin and reproductive tract cancers and to other phenotypes not previously associated with the HLA region (e.g. anemias and acne). We find several non-additive effects, suggesting a more complex landscape of disease-modifying effects throughout the region. Finally, we discover associations between homozygous HLA allele burden and several cancer and other phenotypes, suggesting that peptide presentation spectra as coded for by the HLA region are important in determining disease risk. Our results demonstrate the HLA region’s complexity and richness while underscoring its clinical relevance.

## Introduction

The human leukocyte antigen (HLA) region of the genome is one of the most disease-associated, gene-dense, and polymorphic regions of the human genome^1^. The downstream products of HLA genes generate and present peptides on the cell surface that can be recognized by T cell receptors, making these genes relevant to many disorders of the immune system^2^. Before the advent of high-throughput genome-wide association studies (GWAS), HLA polymorphisms were associated only with HIV, autoimmune disorders, and cancers^3,4^. Since, GWAS has uncovered associations between HLA alleles and common infections^5^, shingles^6^, chronic Hepatitis B^7^, Epstein-Barr virus^8^, and other diseases. The HLA region contains 1.5% of the genes currently in Online Mendelian Inheritance in Man (OMIM, a database of genetic disorders and traits focusing on gene-phenotype relationships) and accounts for roughly 1% (1,827/185,864) of the genome-wide significant single nucleotide polymorphism (SNP) associations in the NHGRI/EBI GWAS catalog^9^, highlighting its importance to disease. Despite being an attractive target for comprehensive GWAS, the HLA region presents many challenges that obscure its roles in disease pathogenesis. Complex linkage disequilibrium (LD) structures, structural variation, closely-related genes, paralogs, segmental duplications, and violations of Hardy-Weinberg equilibrium make high-throughput genotyping methods such as fine-mapping and imputation challenging^10,11^. Chips (e.g. the Illumina MHC SNP Panel^12^) and robust reference panels have been custom-built to study the region^13,14^, but there still exist well-studied alleles in the HLA region whose impact on disease are unknown. Thus, understanding the role of the HLA region is critical to assessing disease risk.

The UK Biobank contains genetic and phenotypic data for over 500,000 individuals, and additionally provides the opportunity to analyze HLA genotypes generated via imputation strategies that address the aforementioned challenges^13^. These data enable association analyses between specific HLA alleles and the rich diversity of phenotypes in the UK Biobank, which are derived from cancer registry, hospital in-patient, primary care, and self-reported questionnaire data. Assessing relationships between HLA alleles and individual-level phenotype data can help pinpoint culprit alleles for disease associations via well-powered conditional analyses. We can additionally identify settings where HLA alleles exhibit non-additive effects on phenotypes, improving our ability to accurately integrate them in disease risk models. Finally, we can test for the effect of overall HLA homozygosity burden on phenotypes, as homozygosity in the HLA region has previously been linked to disease risk^3,4^.

Here, we assessed associations between 156 HLA alleles and 677 binary phenotypes across 337,138 unrelated white British individuals in the UK Biobank and identified 1291 associations at a false discovery rate (FDR) of 5% after Benjamini-Yekutieli (BY) multiple testing correction. The associations spanned 113 HLA alleles (across all 11 Biobank-provided HLA loci) and 128 binary phenotypes. We used Bayesian Model Averaging to reduce the 1291 associations to conditionally independent alleles and uncovered 178 high-confidence (posterior probability >= 80%) associations spanning 50 alleles (across all 11 HLA loci) and 78 binary phenotypes (88 [49%] of which were supported by the literature, **Supplementary Table 1**). We assessed these associations for non-additive effects, i.e. “disease contributions beyond the cumulative effect of individual alleles”^15^, finding 25 associations with significant deviations from additivity (8 [32%] of which were supported by the literature, **Supplementary Table 1, Figure 1**). 9 of these 25 non-additive effects drive intestinal malabsorption and/or celiac disease, but other associations affect a multitude of other autoimmune and endocrine disorders. For example, although HLA-B*27:05 is known to have a strong effect on ankylosing spondylitis and iridicyclitis, and HLA-B*57:01 is likewise known to have a strong effect on psoriasis, we find that these effects are also non-additive. This study associates HLA alleles with a variety of phenotypes in the UK Biobank through association analyses and model selection techniques, providing important insights into disease pathogenesis.

**Figure 1.**
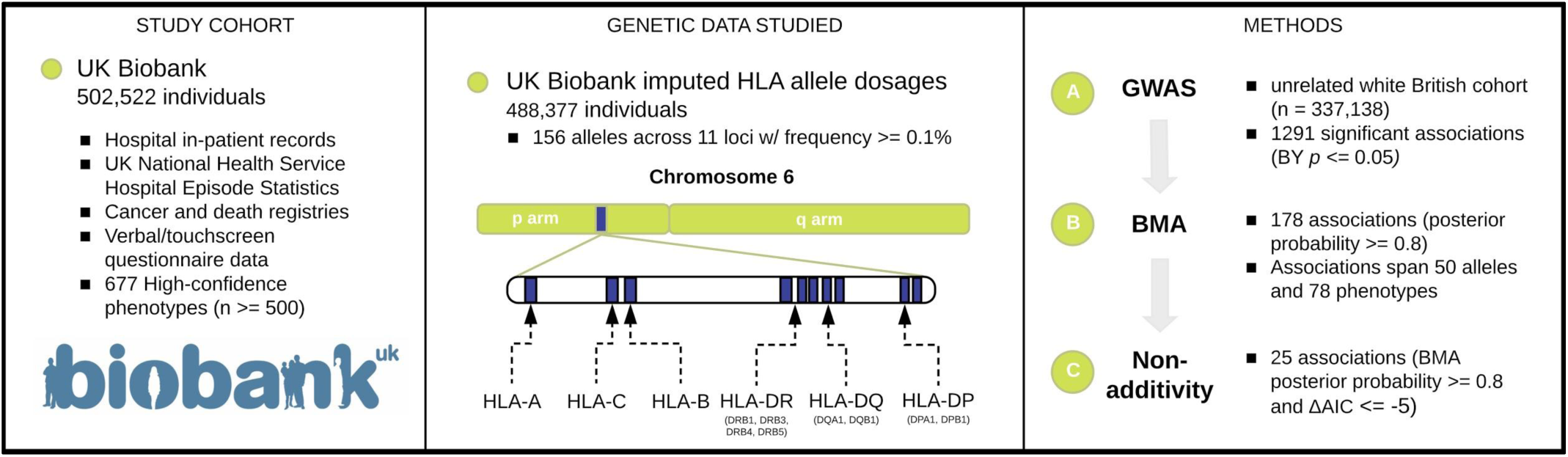
Overview of the study design. We prepared a data set of 156 HLA alleles and 677 medical phenotypes across 337,205 white British individuals in the UK Biobank. We then investigated the effects of these alleles on disease risk.

## Materials and Methods

### Data

For unrelated white British individuals (n = 337,138) in the UK Biobank, four sets of phenotypes were defined: “high-confidence”, “cancer”, “time-to-event”, and “algorithmically-defined”. High-confidence and cancer phenotypes were defined as previously^16^ by mapping ICD-10 codes (from the UK Cancer Register [http://biobank.ctsu.ox.ac.uk/crystal/label.cgi?id=100092], Data-Field 40006 [http://biobank.ctsu.ox.ac.uk/crystal/field.cgi?id=40006], and/or Data-Field 41202 [http://biobank.ctsu.ox.ac.uk/crystal/field.cgi?id=41202]) to self-reported diagnoses (Data-Field 20001 [http://biobank.ctsu.ox.ac.uk/crystal/field.cgi?id=20001] and Data-Field 20002 [http://biobank.ctsu.ox.ac.uk/crystal/field.cgi?id=20002]) from the UK Biobank questionnaire using the FuzzyWuzzy package token set ratio function. Time-to-event phenotypes were derived from First Occurrence of Health Outcomes data as defined by 3-character ICD-10 codes in UK Biobank’s Category 1712. The First Occurrence data-fields were generated by combining: read code information in the primary care data (Category 3000); ICD-9 and ICD-10 codes in hospital inpatient data (Category 2000); ICD-10 codes in death registry records (Field 40001, Field 40002); and self-reported medical condition codes (Field 20002), reported at baseline or subsequent UK Biobank assessment center visits as 3-character ICD-10 codes. Algorithmically-defined outcomes (based on data from Category 42) include phenotypes of select health-related events obtained through algorithmic combinations of coded information from the UK Biobank’s baseline assessment data collection. The data were derived from self-reported medical conditions, operations and medications together with linked data from hospital admissions and death registries. We included phenotypes with at least 500 cases among the white British cohort (**Supplementary Table 2**) and manually deduplicated several phenotypes using the FuzzyWuzzy python package’s ‘partial_ratio’ function on the phenotype names, resulting in the 677 phenotypes used in the analysis.

HLA alleles HLA-A, -B, -C, -DPA1, -DPB1, -DQA1, -DQB1, -DRB1, -DRB3, -DRB4, and -DRB5 were imputed using the HLA*IMP:02 program^13^. HLA-IMP:02 is built on multiple reference panels and a graphical model of the HLA haplotype structure. The UK Biobank provides an imputed dosage file that contains the estimated dosage of 362 HLA alleles across 11 HLA loci from 488,377 individuals (while modeling imputation uncertainty). We included 156 alleles across all 11 loci that had a frequency of 0.1% or greater in our white British cohort (**Supplementary Table 2**). For consistency across models, and because the non-additivity analysis requires integral values for allele dosage, those dosages that were within 0.1 of 0, 1, or 2 for each allele were rounded to integer genotypes, and the remaining nonzero entries were excluded. Erroneous total allele counts post-rounding were excluded.

### Association analysis

We used a generalized linear model (with “Firth-fallback”) as implemented in PLINK v2.00aLM (March 14 2020)^17,18^ to generate association results for the above-described 677 phenotypes and 156 HLA alleles across UK Biobank white British individuals (n = 337,138). Firth-fallback is a hybrid algorithm which normally uses logistic regression but “falls back” to Firth’s bias reduction method^19^, equivalent to penalization of the log-likelihood by the Jeffreys prior, in two cases: (1), one of the cells in the 2×2 (allele count-by-case/control status) contingency table is empty; or (2), logistic regression fails to converge within the usual number of steps. We used age, sex, genotyping array, number and length of copy number variants, and the first ten genotype principal components as covariates in our analysis^20^. We additionally computed an additive model (with the same parameters as PLINK) in R in order to facilitate the subsequent non-additivity analysis. We used the Benjamini-Yekutieli (BY) multiple-testing FDR control method^21^ to adjust the resultant p-values, selecting those associations with BY-corrected p-values less than 0.05 to control FDR at 5%.

### Bayesian Model Averaging

Given the high LD between HLA alleles and the high number of potentially spurious HLA allele-phenotype associations even despite FDR correction, we used a Bayesian Model Averaging (BMA) approach, implemented in the “bma” R package^22^, to prioritize which HLA loci were most likely causal for each phenotype. For a given phenotype, we used BMA to fit a model for each possible combination of significantly-associated alleles. The posterior probabilities of each model are calculated, and the sum of the posterior probabilities of the models in which an allele is included (the “allele posterior probability”) is then a measure of confidence in the allele-phenotype association.

We identified all of the allele-phenotype pairs that had BY-adjusted p-values less than or equal to 0.05 from the additive association analysis for use in BMA. Because BMA is exponential in complexity with respect to the number of alleles analyzed (requiring analysis of 2^n+1^ models for n alleles included in the analysis), we used BMA only on the 20 alleles with the lowest BY-adjusted p-values from the additive association analysis for each phenotype. If there were less than two such alleles for a given phenotype, we did not utilize BMA for that phenotype. Additionally, only the models whose posterior probabilities were within a factor of 1/5 of that of the best model were kept for the final averaging. After applying these filters, we used BMA for 116 phenotypes, with 111 alleles included in at least one model. We used BMA with a binomial link function and error distribution and with age, sex, genotyping array, and the first ten genotype principal components as covariates (number and length of copy number variant covariates were excluded). We focused on allele-phenotype pairs with BMA posterior probabilities >= 0.8 in the subsequent non-additivity analysis.

### Analysis of non-additive genetic effects

To additionally assess whether certain allele-phenotype pairs exhibited non-additive effects on their phenotypes, we used logistic regressions in R (“glm” function, family = “binomial”) and provided HLA allele dosages as factors (i.e., separate terms indicated whether a subject was heterozygous or homozygous for the HLA allele in question). We included age, sex, genotyping array, number and length of copy number variants, and the first ten genotype principal components as covariates (as in the single-allele association analysis). We computed an Akaike Information Criterion (AIC, a measure of goodness-of-fit^23^), comparing the non-additive and additive association models in R for model selection. To identify gene-phenotype associations with suspected departures from additivity, we identified allele-phenotype pairs where the BMA posterior probability was greater than 0.8 and where ΔAIC = AIC_non-additive_ - AIC_additive_ <= −5.

### Homozygosity burden test

To evaluate the burden of homozygous HLA alleles on phenotype, we tabulated the number of homozygous HLA alleles in each individual and fit a logistic model to the standard deviation of this burden in R, with age, sex, genotyping array, genotype missigness, number and length of copy number variants, and the first ten genotype principal components as covariates. We used the Benjamini-Yekutieli (BY) multiple-testing FDR control method^21^ to adjust the resultant *p*-values across all phenotypes, selecting those associations with BY-corrected *p*-values less than 0.05 to control FDR at 5%.

### Literature review

For each allele-phenotype pair resulting from the BMA analysis, we conducted a comprehensive literature review to determine whether the association had been previously found. The HLA allele and the phenotype in question were entered into Google Scholar both with and without quotations, and aliases were checked (e.g. B*08:01, B*0801, B/08/01, and B/0801). Search results were manually inspected to determine associations (**Supplementary Table 1**).

## Results

### Association analysis

To assess the extent to which HLA alleles affect various phenotypes, we conducted an HLA-wide association study across 337,138 white British individuals in the UK Biobank (Methods, **Figure 1**). We used 156 HLA alleles (out of 175 total) that were present at greater than 0.1% minor allele frequency in the cohort and 677 de-duplicated binary phenotypes with more than 500 cases in the white British cohort in the UK Biobank. These binary phenotypes included autoimmune, lymphatic, cardiovascular, dermal, skeletal, tissue, digestive, respiratory, renal, and endocrine disorders as well as many cancers. To control for false discovery, we corrected for multiple testing across the phenotypes using the Benjamini-Yekutieli (BY) procedure at a FDR of 5%. We found 113 alleles that were associated with at least one of 128 binary phenotypes for a total of 1291 associations.

### Bayesian Model Averaging

Given the high degree of LD between HLA alleles, we examined whether the identified associations were conditionally independent. We used Bayesian Model Averaging (BMA) on the top 20 BY-significant alleles for each phenotype that had at least 2 BY-significant allele associations to estimate the posterior effect size estimate, BMA_postOR_. Across 116 test phenotypes and 111 alleles included in at least one analysis, we found 178 associated allele-phenotype pairs among 50 distinct alleles and 78 distinct phenotypes. Of these 178 associations, 88 have been previously documented as associated allele-disease pairs, and 90 were novelly marked by our analysis as high-probability (**Methods, Supplementary Tables 1-2, Figure 2A, Figure 2B**). Of note, 83 allele-phenotype pairs with posterior probability 1 were novel and not previously documented in the literature. These 83 allele-phenotype pairs mostly consist of associations that either feature alleles in LD with known associations or associations between alleles and closely-related phenotypes to known associations. For example, we find an association between **non-melanoma skin cancer and DQB1*03:02**, although the literature reports an association with an allele in LD with DQB1*03:02, DQB1*05:01^24^. Alleles associated to celiac disease, such as **B*08:01**^25^, were found to be associated with **intestinal malabsorption and anemias**, which have close ties to celiac disease^26^. Additionally, alleles associated with Graves’ disease and Hashimoto thyroiditis (such as **B*08:01**^27^, **B*39:06**^28^, **DQB1*06:04**^29^, and **DRB1*03:01**^30^) were found to be associated with **hypo- and hyper-thyroidism** in the UK Biobank and account for 13 of the 83 high-probability associations. Finally, alleles associated with type 1 diabetes (**DQB1*02:01, DQB1*03:02, and DRB1*04:01**) were found to be associated with “**other disorders of pancreatic internal secretion**,” which is biologically related to type 1 diabetes.

**Figure 2.**
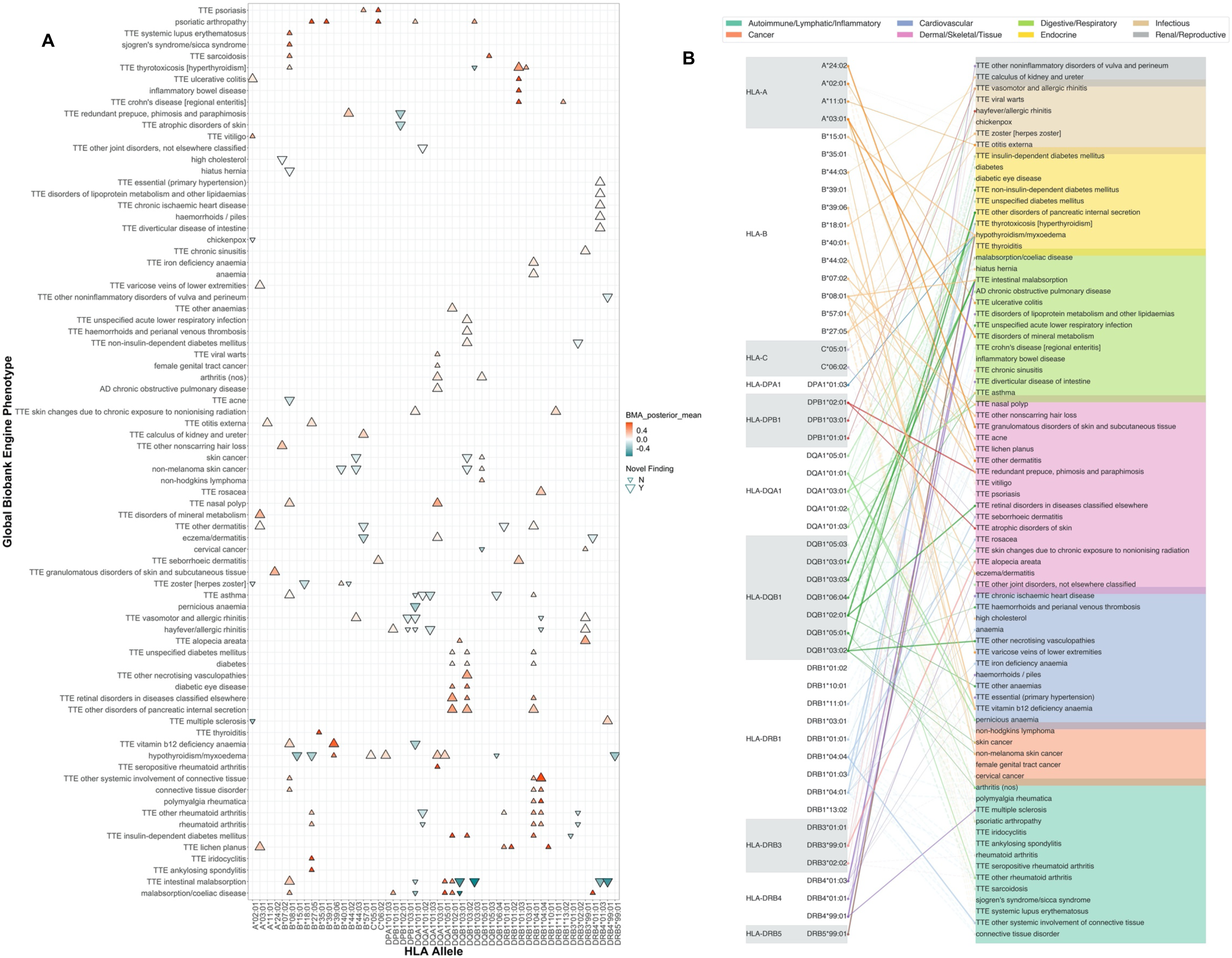
Overview of BMA results. A) Direction and magnitude of HLA allele effects on phenotypes. x-axis indicates HLA allele, y-axis indicates phenotype. Up arrow shows risk effects; down arrow shows protective effects. The effect sizes are coded on a spectrum of red (risk) to blue (protection). A systematic literature search was conducted, yielding a set of previously reported and novel associations (**Supplementary Table 1**). Marker size indicates novelty of discovery - small (previously found) or large (novel). y-axis was clustered by mean effects of alleles on phenotypes. **B) Spider plot showing BMA associations between HLA alleles and binary phenotypes.** The phenotypes were grouped by category as in the legend. A line was drawn between an allele and a phenotype if the allele had a BMA posterior probability ≥ 0.8 for that phenotype. Novel associations (as determined in **Supplementary Table 1**) are shown as solid, opaque lines; previously-found associations are indicated as dashed, faded lines. Line width scales with BMA posterior mean effect size.

Other associations among these 178 include those between HLA alleles and infectious diseases, asthma, systemic autoimmune disorders, and skin cancers (**Supplementary Table 1**). Among the strongest risk effects (posterior probability 1) are associations between: **B*27:05 and ankylosing spondylitis** (p_BY_ = 0, BMA_postOR_ = 9.02, 95% CI = [8.19, 9.94])^31^, **B*27:05 and iridocyclitis**, a type of uveitis (p_BY_ = 2.94 × 10^−239^, BMA_postOR_ = 4.75, 95% CI = [4.32, 5.23])^32^, **B*39:01 and psoriatic arthropathy** (p_BY_ = 1.06 × 10^−8^, BMA_postOR_ = 3.54, 95% CI = [2.51, 5.0])^33^ and **DQA1*03:01 and seropositive rheumatoid arthritis** (p_BY_ = 6.46 × 10^−66^, BMA_postOR_ = 3.16, 95% CI = [2.75, 3.63])^34^. We replicate previously-found strong protective effects of **DQB1*03:01** (p_BY_ = 1.00 × 10^−70^, BMA_postOR_ = 0.42, 95% CI = [0.34, 0.51]) and **DQA1*01:01** (p_BY_ = 2.62 × 10^−51^, BMA_postOR_ = 0.71, 95% CI = [0.61, 0.81])^15^ on **celiac disease** and **malabsorption**. We also discover novel associations to these phenotypes: **DQB1*03:03** (p_BY_ = 2.30 × 10^−22^, BMA_postOR_ = 0.47, 95% CI = [0.36, 0.62]), DQB1*03:01 (p_BY_ = 2.00 × 10^−53^, BMA_postOR_ = 0.42, 95% CI = [0.34, 0.51]), **DRB4*99:01** (p_BY_ = 4.47 × 10^−31^, BMA_postOR_ = 0.52, 95% CI = [0.46, 0.58]), and **DRB4*01:03** (p_BY_ = 1.40 × 10^−71^, BMA_postOR_ = 0.63, 95% CI = [0.49, 0.81]) are all associated. Of note, we find new HLA associations to anemias, with which HLA alleles have not previously been associated with, except for the rare case of aplastic anemia (Latin America, France, Turkey, Brazil, and Barcelona have reported prevalences of only 1.6, 1.5, 0.64, 2.4, and 2.5 per million for the disease respectively)^35^. We find **B*08:01** (p_BY_ = 8.71 × 10^−4^, BMA_postOR_ = 1.2, 95% CI = [1.11, 1.3]) and **B*39:06** (p_BY_ = 7.66 × 10^−3^, BMA_postOR_ = 1.95, 95% CI = [1.49, 2.56]) to be associated with **vitamin B12 deficiency anemia** risk with moderate effects. In contrast, **DQA1*01:01** protects against **vitamin B12 deficiency anemia** (p_BY_ = 8.41 × 10^−7^, BMA_postOR_ = 0.76, 95% CI = [0.68, 0.83]) and **pernicious anemia** as well (p_BY_ = 7.20 × 10^−3^, BMA_postOR_ = 0.73, 95% CI = [0.63, 0.83]).

Several other associations are intriguing. The only previous association between the HLA region and acne is the discovery of the DPB1*04:02 allele itself while HLA-typing a patient with acne vulgaris^36^. Yet, we find a weak protective effect of **B*08:01 on acne** (p_BY_ = 4.21 × 10^−3^, BMA_postOR_ = 0.85, 95% CI = [0.79, 0.91]). This allele is additionally found to be associated with multiple systemic autoimmune disorders, such as **connective tissue disorder** (p_BY_ = 3.45 × 10^−10^, BMA_postOR_ = 1.32, 95% CI = [1.23, 1.42]), **sarcoidosis** (p_BY_ = 2.04 × 10^−33^, BMA_postOR_ = 1.58, 95% CI = [1.36, 1.84]), **Sjogren’s syndrome** (p_BY_ = 1.52 × 10^−15^, BMA_postOR_ = 1.83, 95% CI = [1.61, 2.09]), and **systemic lupus erythematosus** (p_BY_ = 9.50 × 10^−12^, BMA_postOR_ = 1.79, 95% CI = [1.55, 2.06]), suggesting diverse and widespread involvement in various immune and bodily functions (**Supplementary Table 1**). Additionally, we find that **B*57:01** (notably associated with a spike in Alanine aminotransferase levels, among other adverse drug reactions, to the antiretroviral abacavir^37,38^) to be associated with **calculus of kidney and ureter**.

Associations between HLA and cancers are well-established, and here we recover several, including between **DQB1*05:01** and **DRB3*99:01** and **cervical cancer** and **DQA1*03:01** and **female genital tract cancer**. Furthermore, we find **DQB1*05:01** to be associated with **skin cancer, non-melanoma skin cancer**, and **non-hodgkin’s lymphoma**. In addition, we identify several novel associations between skin cancer, non-melanoma skin cancer, and HLA alleles. Statistics on all cancer associations (as well as their novelty) are described in **Table 1**. Taken together, by applying BMA, we recover known associations as well as identify novel disease associations with HLA alleles. These additive associations could add to elucidation of disease risk across a multitude of complex traits.

**Table 1.**
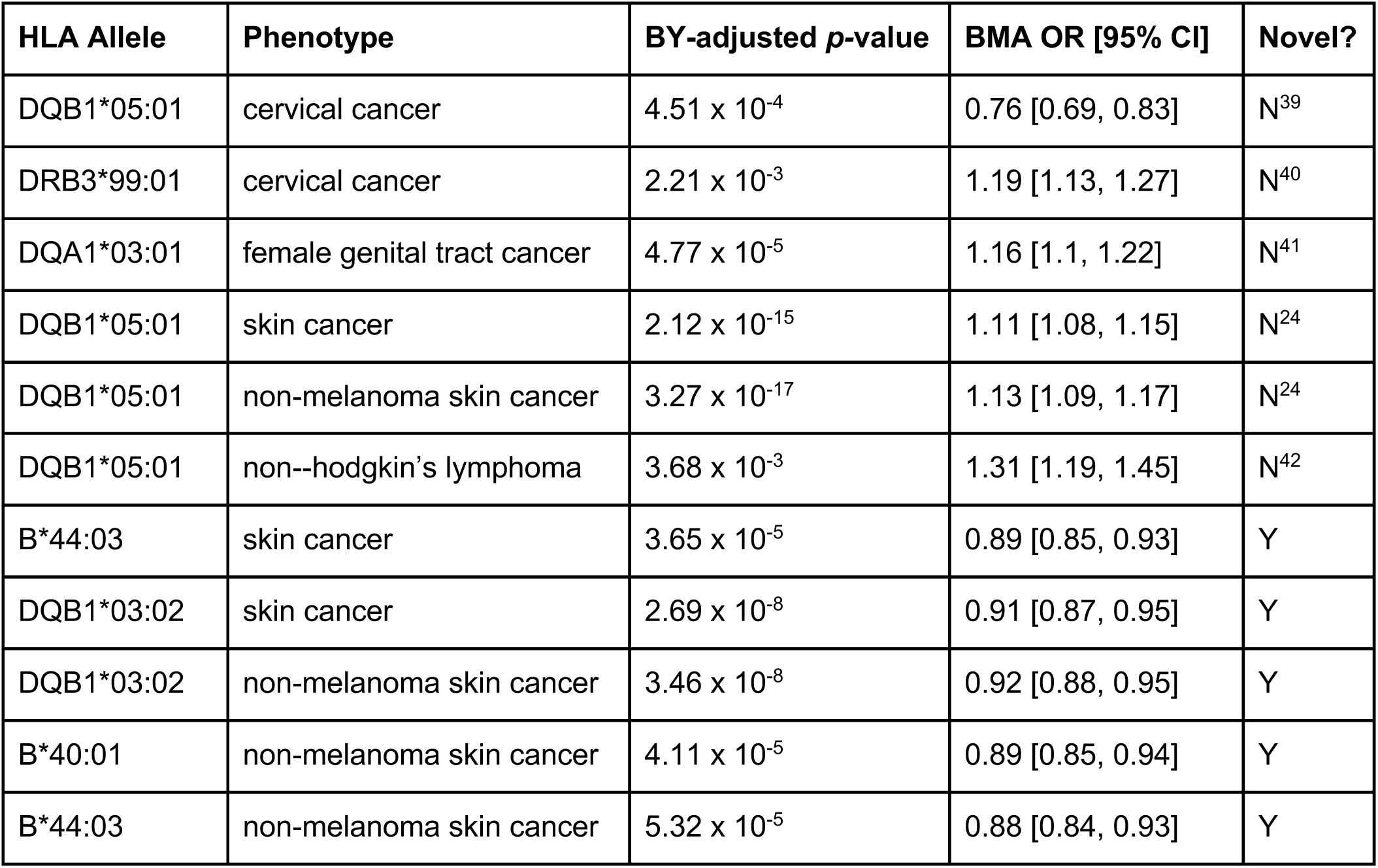
Cancer associations found by BMA (posterior probability >= 0.8). For each cancer association (“HLA Allele” and “Phenotype”), the table provides: “BY-adjusted *p-*value”: Benjamini-Yekutieli-adjusted *p*-value from the univariate association analysis (*p-*values were adjusted across all phenotypes); “BMA OR [95% CI]”: odds ratio from the BMA analysis alongside 95% confidence intervals; “Novel?”: whether or not each BMA-significant cancer association is novel. If not novel, citation is provided.

### Analysis of non-additive genetic effects

Several analyses have found evidence of non-additive effects within the HLA region of the genome^15^. Thus, we examined whether non-additive effects among the 178 associations identified in the BMA analysis contribute to disease risk. Where ΔAIC was defined as the Akaike Information Criterion goodness-of-fit^23^ difference between the additive (standard GWAS using numerical values) model and the non-additive/”genotypic” model (in which rounded genotype dosage is considered as a factor), we selected those associations where ΔAIC = AIC_non-additive_ - AIC_additive_ <= -5, i.e. where the genotypic model fit was significantly better than that of the additive model.

We found 25 associations which met this ΔAIC criterion, of which 8 were previously described^15^ (**Figure 3**). There is an enrichment for malabsorption (9) thyroid (4), and autoimmune (4) phenotypes, but psoriatic (3), asthma (2), and generic diabetes (2) phenotypes also feature multiple non-additive associations. We define heterozygote odds ratio (OR) to be the OR associated with Dosage 1 and homozygote OR to be the OR associated with Dosage 2, as in **Figure 3**. We replicate a known non-additive effect^15^ of **C*06:02** on **psoriasis** (additive OR 2.79, 95% CI [2.69, 2.89]; heterozygote OR 3.19, 95% CI [3.05, 3.33]; homozygote OR 4.35, 95% CI [3.79, 4.99]). In addition, we identify multiple novel associations, such as **B*27:05** and **ankylosing spondylitis** (additive OR 8.93, 95% CI [8.13, 9.81]; heterozygote OR 12.24, 95% CI [10.94, 13.68]; homozygote OR 14.11, 95% CI [8.24, 24.18]); low homozygote counts result in a large CI for this homozygote OR. In relation to the association between **DRB5*99:01** and **hypothyroidism/myxedema** (additive OR 1.22, 95% CI [1.19, 1.26]; heterozygote OR 1.44, 95% CI [1.28, 1.62]; homozygote OR 1.72, 95% CI [1.53, 1.93]), the heterozygote and homozygote CIs overlap with each other but not with the additive CI. We see a significantly higher homozygote OR for **B*57:01** and **psoriasis** (additive OR 3.07, 95% CI [2.93, 3.23]; heterozygote OR 3.22, 95% CI [3.06, 3.39]; homozygote OR 5.42, 95% CI [4.14, 7.09]), representing another type of departure from additivity. Other non-additive associations have CIs that all overlap with each other, e.g. **B*27:05** and **iridocyclitis** (additive OR 4.83, 95% CI [4.4, 5.29]; heterozygote OR 5.5, 95% CI [4.96, 6.09]; homozygote OR 8.05, 95% CI [4.8, 13.51]). The effect of **DRB4*01:01 on celiac disease** (additive OR 1.71, 95% CI [1.56, 1.87]; heterozygote OR 1.92, 95% CI [1.74, 2.12]; homozygote OR 0.62, 95% CI [0.3, 1.31]) has a homozygote effect estimate somewhat opposite that of the heterozygote and additive effect estimates, albeit the CI of the homozygote OR crossing 0 (**Figure 3**). This could indicate a recessive effect.

**Figure 3.**
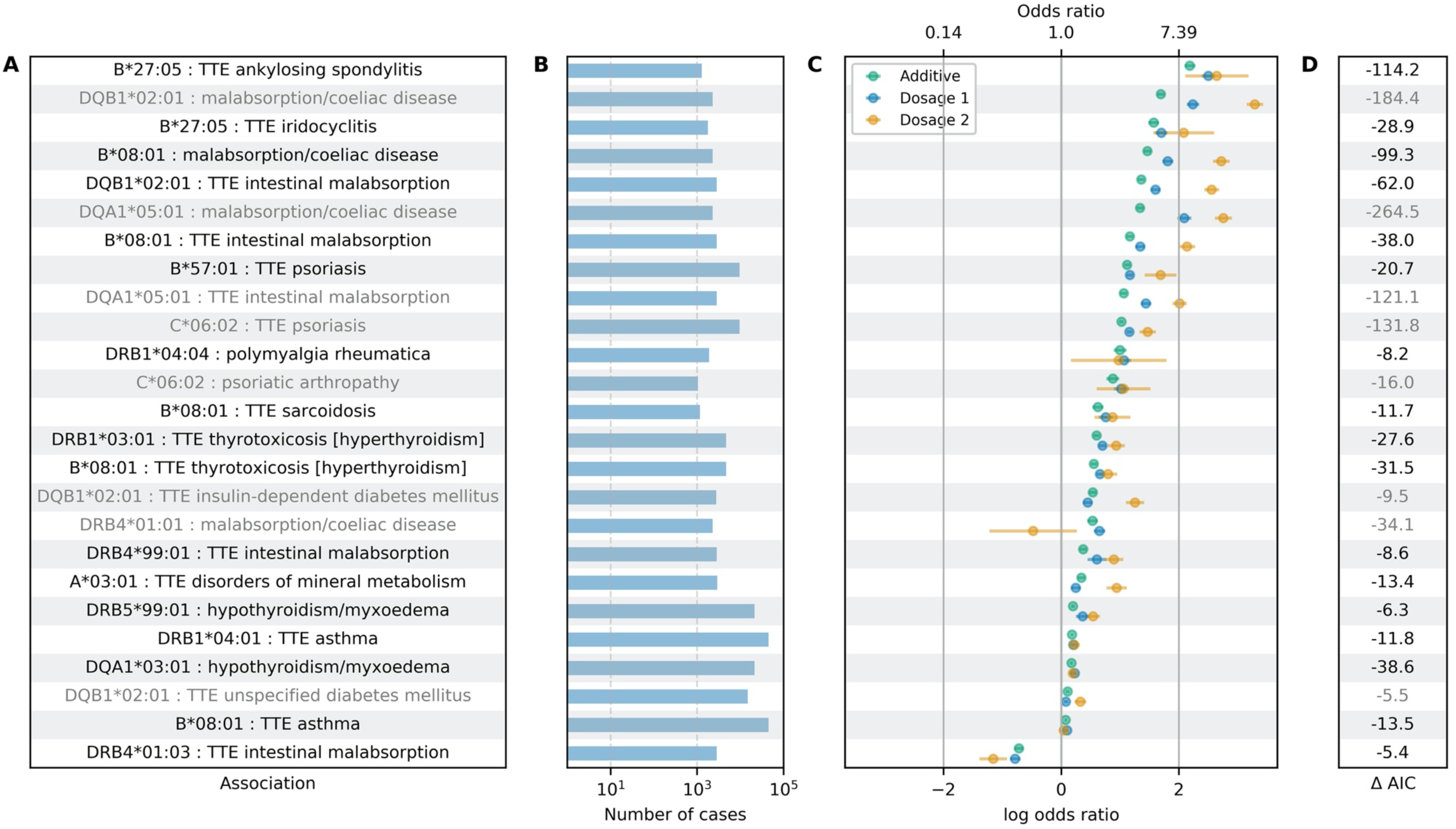
Non-additive dosage model associations. (A) Allele-phenotype pair. Light grey text represents a previously-found non-additive effect (**Supplementary Table 1**). **(B) Number of UK Biobank cases for the phenotype** x-axis is log_10_ scaled. **(C) Odds ratio and log odds ratio for additive (green) and genotypic model (blue, dosage 1 and yellow, dosage 2).** Graphical measure of model fit. **(D) ΔAIC (AIC additive model - AIC genotype model).** A more negative value represents a larger departure from additivity.

12 previously-known moderate non-additive associations^15^ were not replicated. In our analyses, these range in ΔAIC from -3.368 to +1.99 (**Supplementary Table 2**). Overall, the underlying biology of these non-additive associations could be important to disease pathogenesis.

### Homozygosity burden test

Homozygosity of HLA alleles is purported to have adverse effects on disease risk, especially with respect to cancer and infectious diseases, because of the decrease in peptide presentation spectra on the cell surface^43^. An individual with two distinct HLA alleles at a locus will have a larger spectrum of presentable peptides, and in turn, the probability that this larger spectrum includes disease-relevant peptides is also higher. To test for the effect of homozygosity on disease risk, we used a logistic regression with the standard deviation of the number of homozygous HLA alleles per individual and the 677 phenotypes as input. As hypothesized, the BY-significant associations (**Table 2**) are enriched for cancers, but also include asthma, colitis, autoimmune, and various allergic and skin conditions. All but the associations to rhinitis and asthma are weak risk effects. The results thus indicate that HLA allele homozygosity may introduce a weak baseline risk towards certain cancers and autoimmune diseases, and weak baseline protection towards others.

**Table 2.**
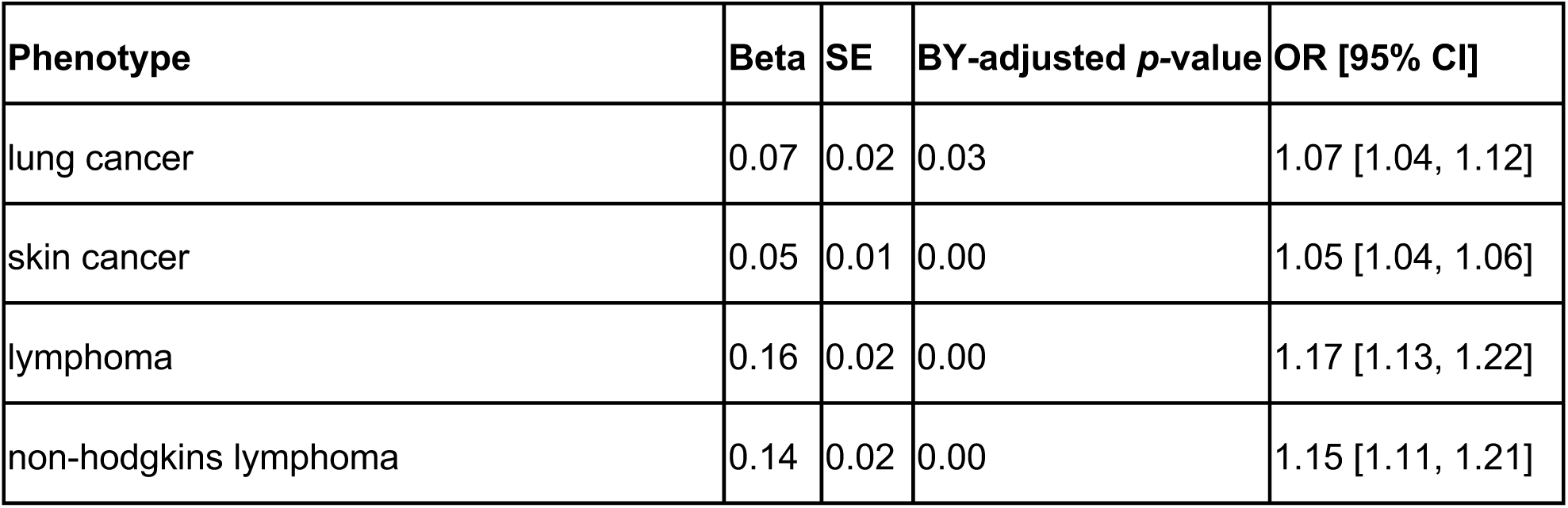
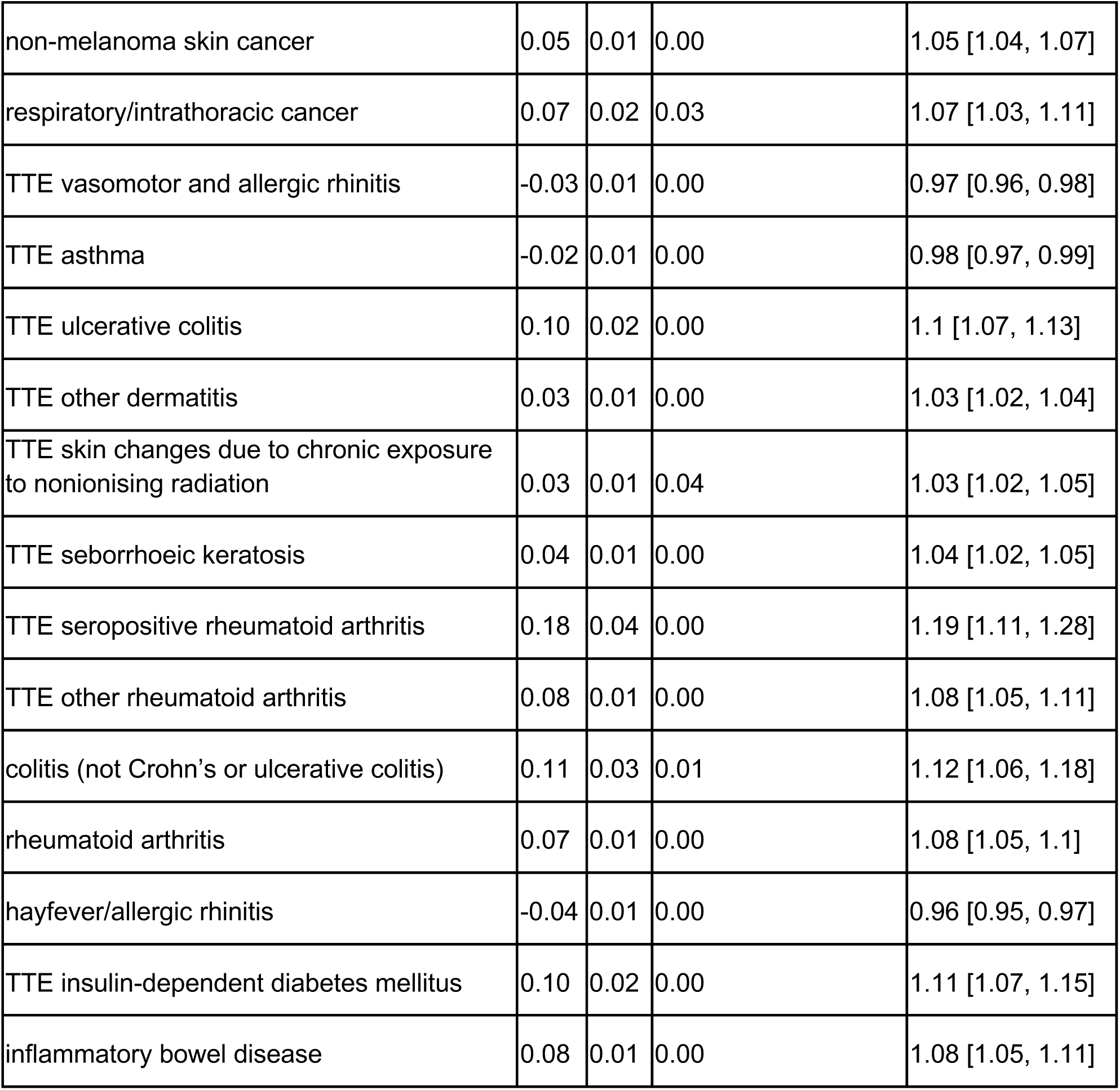
Associations to homozygosity burden. “Beta”: log odds ratio (OR) per standard deviation of number of homozygous HLA alleles per individual (SD = 1.84). “SE”: standard error of the log odds ratio as provided in column “Beta”. “BY-adjusted *p-*value”: Benjamini-Yekutieli-adjusted *p*-value from association analysis between standard deviation of number of homozygous HLA alleles per individual and phenotype. *p*-values were BY-adjusted across all phenotypes. “OR [95% CI]”: Odds ratios corresponding to the “Beta” column and their 95% confidence intervals.

| Phenotype | Beta | SE | BY-adjusted *p*-value | OR [95% CI] |
| --- | --- | --- | --- | --- |
| lung cancer | 0.07 | 0.02 | 0.03 | 1.07 [1.04, 1.12] |
| skin cancer | 0.05 | 0.01 | 0.00 | 1.05 [1.04, 1.06] |
| lymphoma | 0.16 | 0.02 | 0.00 | 1.17 [1.13, 1.22] |
| non-hodgkins lymphoma | 0.14 | 0.02 | 0.00 | 1.15 [1.11, 1.21] |
| non-melanoma skin cancer | 0.05 | 0.01 | 0.00 | 1.05 [1.04, 1.07] |
| respiratory/intrathoracic cancer | 0.07 | 0.02 | 0.03 | 1.07 [1.03, 1.11] |
| TTE vasomotor and allergic rhinitis | -0.03 | 0.01 | 0.00 | 0.97 [0.96, 0.98] |
| TTE asthma | -0.02 | 0.01 | 0.00 | 0.98 [0.97, 0.99] |
| TTE ulcerative colitis | 0.10 | 0.02 | 0.00 | 1.1 [1.07, 1.13] |
| TTE other dermatitis | 0.03 | 0.01 | 0.00 | 1.03 [1.02, 1.04] |
| TTE skin changes due to chronic exposure to nonionising radiation | 0.03 | 0.01 | 0.04 | 1.03 [1.02, 1.05] |
| TTE seborrhoeic keratosis | 0.04 | 0.01 | 0.00 | 1.04 [1.02, 1.05] |
| TTE seropositive rheumatoid arthritis | 0.18 | 0.04 | 0.00 | 1.19 [1.11, 1.28] |
| TTE other rheumatoid arthritis | 0.08 | 0.01 | 0.00 | 1.08 [1.05, 1.11] |
| colitis (not Crohn's or ulcerative colitis) | 0.11 | 0.03 | 0.01 | 1.12 [1.06, 1.18] |
| rheumatoid arthritis | 0.07 | 0.01 | 0.00 | 1.08 [1.05, 1.1] |
| hayfever/allergic rhinitis | -0.04 | 0.01 | 0.00 | 0.96 [0.95, 0.97] |
| TTE insulin-dependent diabetes mellitus | 0.10 | 0.02 | 0.00 | 1.11 [1.07, 1.15] |
| inflammatory bowel disease | 0.08 | 0.01 | 0.00 | 1.08 [1.05, 1.11] |

## Discussion

This study is a comprehensive overview of associations between 156 HLA alleles and 677 phenotypes in 337,138 individuals in the UK Biobank. Using single-allele association analysis and subsequent BMA, we replicate 88 known associations between HLA alleles and binary phenotypes (including cancers) and discover 90 novel associations. Many of the novel associations feature phenotypes close to known associations (e.g. intestinal malabsorption and celiac disease, hypo- and hyperthyroidism), but others are completely novel (e.g. anemias and acne). For example, we find that DRB1*04:04 is associated with a family of autoimmune disorders such as rheumatoid arthritis, polymyalgia rheumatica, and connective tissue disorder, replicating signals found previously in the literature. We add to the body of known HLA associations strong, novel effects such as that of B*39:06 on vitamin B12 deficiency anemia. Finally, we find several novel associations to skin and reproductive tract cancers, highlighting the clinical importance of the HLA region.

These genotype-phenotype associations implicate the HLA region of the genome in contributions to many diverse phenotypes and suggest avenues for uncovering relevant, novel biology and developing therapeutics; however, there are many opportunities for future studies to extend this analysis. One would be to translate HLA alleles into their respective peptide products and analyze whether individual amino acids, or “pockets,” are associated with disease. Amino acids influence the binding properties of the HLA peptide binding groove; then, the source of the effect that is attributed to the presence of an HLA allele could ostensibly be the presence of a particular amino acid motif (which can be shared by multiple HLA alleles)^3^. Also, given that BMA is exponential in complexity with respect to the number of alleles analyzed, it is computationally infeasible to analyze all BY-significant alleles together for certain phenotypes. As such, it is likely that the 178 allele-phenotype associations we find are actually conservative, highlighting the impact of the HLA on disease risk. It could also be valuable to incorporate significantly-associated HLA region non-coding SNPs into the BMA analyses to fine-map association signal at higher resolution. Additionally, the non-additivity analysis may be underpowered to determine homozygote ORs in some cases where those genotypes are low in number. We do not find any non-additive effects consistent with a recessive-only model, i.e. where the heterozygote OR overlaps 1; this is perhaps because homozygosity for an HLA allele translates to a smaller spectrum of peptides displayed on the cell surface and recognized by the immune system. It is also unclear whether or not these non-additive effects can be classified as truly dominant, recessive, or co-dominant effects without additional formal model comparisons. These observations are further supported by our analysis of the effect of burden of homozygous HLA alleles, which suggest that peptide presentation spectra are increasingly important in cancers, infectious diseases, and autoimmune disorders.

HLA alleles are inherited as haplotypes that group several alleles in LD with each other. Better HLA typing could enable accurate haplotype frequency estimation and subsequent haplotype-based tests that enable testing of other targeted hypotheses of HLA diversity. Interactions between HLA alleles are also not studied here and would be a logical extension of this work; examples in which interaction effects are responsible for disease risk include the effect of an *HLA-B/ERAP1* interaction on ankylosing spondylitis^44^ and *HLA-C/ERAP1* on psoriasis^45^. One of the functions of *ERAP1* is to “trim” peptides before they are presented on HLA class I proteins. These studies therefore suggest that the risk effect mediated by certain HLA alleles also depends on peptide pre-processing. As better reference panels, superior imputation techniques, and larger population biobanks with more sequence data are developed, the HLA region will continue to grow in importance in human genetics, and in the future, as effective therapeutic hypotheses are generated, in clinical settings.

## Author Contributions

M.A.R. conceived and designed the study. G.R.V., J.E.O., and C.D. analyzed the data. G.R.V., Y.T., and J.M.J. provided quality control analysis. The manuscript was written by G.R.V. and M.A.R. and revised by all the co-authors. All co-authors have approved of the final version of the manuscript.

## Acknowledgments and Funding

This research has been conducted using the UK Biobank Resource under Application Number 24983, “Generating effective therapeutic hypotheses from genomic and hospital linkage data” (http://www.ukbiobank.ac.uk/wp-content/uploads/2017/06/24983-Dr-Manuel-Rivas.pdf). We thank all of the participants in the UK Biobank study. This work was supported by the National Human Genome Research Institute (NHGRI) of the National Institutes of Health (NIH) under award R01HG010140. The content is solely the responsibility of the authors and does not necessarily represent the official views of the National Institutes of Health. Some of the computing for this project was performed on the Sherlock cluster. We would like to thank Stanford University and the Stanford Research Computing Center for providing computational resources and support that contributed to these research results. G.R.V. is supported by the National Library of Medicine (NLM) T15 Continuing Education Training Grant. J.O. is supported by the National Science Foundation Graduate Research Fellowship under Grant No. DGE-1656518 and a Stanford Graduate Fellowship. Y.T. is supported by a Funai Overseas Scholarship from the Funai Foundation for Information Technology and the Stanford University School of Medicine. J.M.J. is funded by grant NNF17OC0025806 from the Novo Nordisk Foundation and the Stanford Bio-X Program. M.A.R. is supported by Stanford University and a National Institute of Health center for Multi- and Trans-ethnic Mapping of Mendelian and Complex Diseases grant (5U01 HG009080).

## Supplementary Materials

**Supplementary Table 1. Results of systematic literature review.** Allele-phenotype pairs were marked as “novel” if not found in the literature and “non-additive” if the allele was found to have a non-additive effect on the phenotype. PMIDs and URLs for all references were provided. https://bit.ly/hla_sup_1

**Supplementary Table 2. Significant BMA results table.** Results from PLINK additive association analysis, BMA analysis, and non-additivity analysis for each allele-phenotype pair crossing BMA posterior probability 0.8. Number of cases per phenotype and allele frequencies per allele in the UK Biobank white British cohort also provided. https://bit.ly/hla_sup_2

